# The magnitude of cryptic insect diversity in one tropical rainforest

**DOI:** 10.1101/2025.02.28.640797

**Authors:** Yves Basset, Greg P. A. Lamarre, David A. Donoso, Daniel Souto-Vilarós, Filonila Perez, Ricardo Bobadilla, Yacksecari Lopez, José Alejandro Ramírez Silva, Héctor Barrios

## Abstract

1. Cryptic species represent significant entities in conservation planning and are common among tropical insects. Many studies focusing on cryptic species have been restricted to a narrow taxon and it is not clear whether some insect clades or assemblages may have a greater propensity for cryptic speciation than others. Here, we contrast cryptic diversity among 22 insect assemblages surveyed on Barro Colorado Island (BCI), a tropical rainforest in Panama.
2. Cryptic species were defined as externally indistinguishable from their sister species but with a different barcode index number. We detected 214 cryptic species out of a total of 2,006 species studied (10.6%). The percentage of cryptic species varied greatly among assemblages (0-19%), with half of assemblages devoid of cryptic species, and the highest proportions of cryptics in Pieridae and Formicidae.
3. The percentage of cryptic species was weakly influenced by phylogeny and it was best explained by the local number of species and an index of taxonomic knowledge. Cryptic species were distinguished by an average distance divergence in sequence of 9%, but their ecological characteristics remain unclear.
4. Asymptotic estimates indicated that 11.4% of species studied on BCI may be cryptic. Since many of the species-rich insect assemblages occurring on BCI were not considered in this study, this estimate probably represents a lower bound to the true cryptic insect richness on BCI.
5. This significant contribution to biodiversity should not be ignored and, because of apparent low population levels, may be particularly vulnerable to land use and climate changes.

## INTRODUCTION

Cryptic species represent a field of investigation critical for conservation, bioprospecting and biological control (Beheregaray & Caccone, 2007; Bickford et al., 2007; Struck et al., 2018; Cheng *et al*., 2024). The ecology, growth rate or population dynamics of cryptic species may be different from those of non-cryptic species (Bickford et al., 2007). This issue has significant implications. First, biodiversity estimates of certain taxonomic groups may be largely underestimated (Pfenninger & Schwenk, 2007). Second, ecological interactions and networks may be obscured or difficult to disentangle, unless using molecular methods (Ficetola & Taberlet, 2023). Cryptic diversity may also represent an important factor influencing future conservation decisions and efforts (Pfenninger & Schwenk, 2007; Trontelj & Fišer, 2009). Last, misidentification of medically and economically important species in cryptic complexes, such as cryptic pathogens, parasites and invasive species, can also be detrimental to human well-being (Bickford et al., 2007; Pfenninger & Schwenk, 2007).

Cryptic species may be defined with different criteria, depending on whether the perspective is evolutionary, taxonomic or genomic. A classical definition may be “two or more distinct species classified as a single species because they are at least superficially morphologically indistinguishable” (Bickford et al., 2007; Hawksworth, 2012). Other definitions emphasize the role of traditional taxonomy in defining cryptic species (Schönrogge et al., 2002; Karanovic et al., 2016). Cryptic species can also be defined based on their low levels of phenotypic (morphological) disparity relative to their degree of genetic differentiation (Struck et al., 2018). For a historical review of the concept of cryptic species, see Cheng et al. (2024).

Cryptic species are common in the animal kingdom (Bickford et al., 2007; Pfenninger & Schwenk, 2007). For example, they have been confirmed for fungi, sponges, rotifers, trematodes, fishes and insects (e.g., Schönrogge et al., 2002; Nadler & De León, 2011; Hawksworth, 2012). Pfenninger & Schwenk (2007) argued that cryptic species are evenly distributed among major animal taxa and biogeographical regions. However, this view was disputed by Trontelj and Fišer (2009) who stressed the multiple origins and mechanisms generating cryptic species diversity. In support of this argument, extreme environments may support many cryptic species due to selection reducing or eliminating morphological change accompanying speciation (Bickford et al., 2007).

For insects, Stork (2018) noted that unbiased studies of well-researched insect faunas report that 1-2% of species may be truly cryptic. However, a sample of 20 insect studies including at least 10 species (Table S1) reveals that this percentage varies a lot and may be far higher, especially in regional studies (Table S1; see also compilation at the genus level in Li and Wiens, 2023). Li and Wiens (2023) even suggested that each morphologically-based insect species may contain up to 3.1 cryptic species. Cryptic species may be particularly common among tropical insects (Table S1), as exemplified by the hesperiid genus *Astraptes* (Hebert et al., 2004). Although a lot of studies reported cryptic species, their discussion was usually restricted to a narrow taxon (e.g., for insects: Hebert et al., 2004; Smith et al., 2006; Burns et al., 2008; Hendrichs et al., 2015; see Table S1). Multi-taxic studies encompassing insect taxa phylogenetically distant are few (Dincă et al., 2015; Table S1). Hence, it is not clear whether some insect clades or groups may have a greater propensity for cryptic speciation than others (Bickford et al., 2007). In this study, we contrast cryptic diversity among 22 insect assemblages surveyed in one tropical rainforest. The comparison of different assemblages at a single location may reduce confounding factors, such as, for example, different speciation rates in different habitats (Ribera et al., 2001).

Three main hypotheses may explain the occurrence of cryptic species (and see Cheng et al., 2024 for a thorough review): (a) recent divergence (i.e., cryptic species have diverged recently, and morphological differences are not evident); (b) phylogenetic niche conservatism (i.e., niche evolution and morphological differentiation across descendant species are constrained by selection); and (c) morphological convergence (i.e., cryptic species are due to morphological convergence shaped by similar selection pressures; Fišer et al., 2018). Hence, related to (a) the historical age of taxa may be important and evolutionary recent taxa may include high cryptic diversity (Bickford et al., 2007; Struck et al., 2018). Related to (b) sister relationships and the phylogeny of species per se may also be significant (Fišer et al., 2018). Related to (c) taxa with highly specific dietary requirements, such as herbivorous species, may also include many cryptic species (Hebert et al., 2004), as well as taxa with poor dispersal abilities (Jörger & Schrödl, 2013). Other factors explaining contrasting levels of cryptic species among taxa may be related to the overall species pool and the sampling effort to discover cryptic species (Janzen et al., 2012). Thus, we expect a mixture of phylogenetic and ecological effects to explain differing cryptic diversity among insect taxa.

Distinguishing cryptic species on morphology alone may be impossible if a strict definition is enforced or rather subjective for an untrained observer. The second point has merit to be considered. The number of expert taxonomists able to recognize distinct species but very close morphologically in a particular insect group is declining very fast (Leather, 2009). Hence most biologists consulting an entomology collection are likely to be relatively “untrained” eyes. A good alternative to distinguish “true cryptic” species are molecular methods (Belfiore et al., 2003; Bickford et al. 2007; Murray et al., 2008; Nadler & De León, 2011; Struck, et al., 2018). Delineation of insect species is relatively straightforward by sequencing the cytochrome c oxidase subunit I (COI) gene (DNA barcodes: Hebert et al., 2003; Hajibabaei et al., 2006). Interim taxonomic nomenclature is possible by considering COI-based clusters and Barcode Index Number (BIN: Ratnasingham & Hebert, 2013) and has been often used with tropical insects (Hebert et al., 2004; Smith et al., 2006; Burns et al., 2008). Although some controversies exist about the use of BINs to delineate species (Honeycutt, 2021; Cheng et al., 2023; Meier *et al*., 2022), this concept remains the most operational and efficient to date when studying diverse communities in terms of both phylogeny and species richness, such as insects in tropical rainforests (Miller, 2007; Basset et al., 2023).

In 2009, the Arthropod Program of the Smithsonian Tropical Research Institute started long-term monitoring of different insect taxa at Barro Colorado Island (BCI; Lamarre et al., 2020). Since many species collected within this program have been sequenced, this provides a unique opportunity to probe the extend of cryptic diversity among 22 different insect assemblages recruiting from different orders, all originating from a single locality situated in one tropical rainforest.

In this context, we ask the following questions:

1. Does the proportion of cryptic species vary among insect assemblages within the rainforest of BCI? Which of the taxa studied supports a proportionally high share of cryptic species?
2. Accounting for phylogeny, can ecological variables available for the insect assemblages studied explain the proportion of cryptic species among these assemblages?
3. What may be the salient characteristics of the species documented as cryptic?
4. What may be the magnitude of cryptic species in the tropical rainforest of BCI?

## MATERIAL AND METHODS

### Study site and insect data

Insect data were obtained from Barro Colorado Island in Panama (BCI; 9.15◦N, 79.85◦W; 120–160 m asl). BCI receives an average annual rainfall of 2,662 mm, with an annual average daily maximum and minimum air temperatures of 31.0 ◦C and 23.6 ◦C, respectively (https://biogeodb.stri.si.edu/physical_monitoring/research/barrocolorado). The 1,542ha protected island is covered with evergreen wet lowland rainforest and was created around 1910, when the Chagres River was dammed to fill the Panama Canal. Although BCI is physically isolated by the Gatun Lake, physical barriers are weak as the minimal distance to mainland is <500m. Insects were collected within and near the 50ha ForestGEO plot, which is thoroughly described in AndersonTeixeira et al. (2015). Taxonomic knowledge of BCI insects is reasonably good, at least for a tropical location and for well-studied taxa such as butterflies (e.g., Basset et al., 2015). All taxa monitored by the STRI Arthropod Program on BCI since 2009 (Lamarre et al., 2020) are considered in this contribution and are listed in Table 1. Collecting protocols represent standard and reproducible entomological methods and are all detailed in Basset et al. (2013, 2023). Insects were identified by building reference collections improved by expert taxonomists (Table 1). Voucher specimens were deposited in the collection of the STRI Arthropod Program at the Smithsonian Tropical Research Institute in Panama. Monitoring data are stored into the STRI Arthropod database at https://fgeoarthropods.si.edu/. The COI gene was sequenced at the Canadian Centre for DNA Barcoding (Hebert et al., 2003) for representative specimens (number of sequences detailed in Table 1). BINs were used to differentiate species (Hebert et al., 2004) with interim taxonomy when necessary (Ratnasingham & Hebert, 2013). This subsequent sequencing protocol helped to recognize cryptic species indistinguishable morphologically (see below) and to improve reference collections. Molecular data are stored into different public projects of the Barcode of Life Data System (BOLD), as indicated in Table 1, and gradually released into GenBank.

**TABLE 1.** Insect assemblages studied at BCI. Data refer to the period 2009-2022. Herbivores: SS = sap-sucking; LF = leaf-chewing.

| Assemblage | Order | Guild | No.<br>individuals | No.<br>species | No.<br>sequences | BOLD<br>project | Main protocol(s) | Expert(s) |
| --- | --- | --- | --- | --- | --- | --- | --- | --- |
| Kalotermitidae | Isoptera | Saproxylophagous | 8786 | 6 | 28 | BCIIS | Transect & light trap | Rudolf H. Scheffrahn, Jan Krecek |
| Rhinotermitidae | Isoptera | Saproxylophagous | 7457 | 4 | 29 | BCIIS | Transect & light trap | Rudolf H. Scheffrahn, Jan Krecek |
| Termitidae (1) | Isoptera | Saproxylophagous | 22059 | 18 | 130 | BCIIS | Transect & light trap | Rudolf H. Scheffrahn, Jan Krecek |
| Flatidae | Hemiptera | Herbivore (SS) | 4721 | 27 | 113 | BCIFL | Light trap | Edwin Dominguez, Steve W. Wilson |
| Reduviidae | Hemiptera | Predator | 3171 | 86 | 125 | BCIRE | Light trap | Dimitri Forero |
| Passalidae | Coleoptera | Saproxylophagous | 1497 | 22 | 56 | BCIPA | Light trap | Jack C. Schuster |
| Platypodinae | Coleoptera | Saproxylophagous | 1700 | 16 | 57 | BCIPL | Light trap | Héctor Barrios |
| Dynastinae | Coleoptera | Saproxylophagous | 4721 | 31 | 105 | BCIDY | Light trap | Héctor Barrios |
| Hesperiidae | Lepidoptera | Herbivore (LF) | 2078 | 171 | 431 | BCIBT | Pollard walk | Andrew D. Warren |
| Papilionidae | Lepidoptera | Herbivore (LF) | 715 | 16 | 52 | BCIBT | Pollard walk | Robert B. Srygley |
| Pieridae | Lepidoptera | Herbivore (LF) | 5609 | 21 | 62 | BCIBT | Pollard walk | Robert B. Srygley |
| Nymphalidae | Lepidoptera | Herbivore (LF) | 8053 | 133 | 442 | BCIBT | Pollard walk | Robert B. Srygley |
| Lycaenidae | Lepidoptera | Herbivore (LF) | 475 | 56 | 171 | BCIBT | Pollard walk | Robert K. Robbins |
| Riodinidae | Lepidoptera | Herbivore (LF) | 9581 | 56 | 142 | BCIBT | Pollard walk | Robert B. Srygley |
| Geometridae | Lepidoptera | Herbivore (LF) | 22549 | 264 | 1023 | BCIBT | Light trap | Axel Hausmann, Gunnar Brehm, Antoine Lévêque |
| Arctiinae | Lepidoptera | Herbivore (LF) | 29859 | 193 | 874 | BCIAR | Light trap | Michel Laguerre, Benoit Vincent |
| Crambidae | Lepidoptera | Herbivore (LF) | 33873 | 340 | 859 | BCIPY | Light trap | Alma Solis |
| Pyalidae | Lepidoptera | Herbivore (LF) | 15108 | 107 | 88 | BCIPY | Light trap | Alma Solis |
| Saturniidae | Lepidoptera | Herbivore (LF) | 2378 | 45 | 179 | BCISA | Light trap | Rodolphe Rougerie |
| Formicidae | Hymenoptera | Ant | 70580 | 356 | 2447 | BCIFO | Winkler & light trap | David A. Donoso |
| Halictidae | Hymenoptera | Pollinator | 9794 | 6 | 14 | BCIBE | Light trap | David W. Roubik |
| Euglossini | Hymenoptera | Pollinator | 61648 | 32 | 117 | BCIBE | Cineole baits | David W. Roubik |
| All | *** | *** | 326412 | 2006 | 7544 | ABCI | *** | *** |

(1) Without Apicotermiteini

### Categories of cryptic species

In this contribution we take a pragmatic approach to evaluate the proportion of cryptic species in each of the insect assemblage studied. We assigned species to the six following categories, which we consider to be mutually exclusive. We limited the species pool (and comparison between species) to species collected on BCI.

1. **Cryptic identified**: species identified with a binomial, with a distinct BIN (or sequence), indistinguishable from another species when considering external morphology.
2. **Cryptic not identified**: species not identified (possibly identified to genus), with a distinct BIN (or sequence), indistinguishable from another species when considering external morphology.
3. **Non-cryptic identified**: species identified with a binomial, with a distinct BIN (or sequence).
4. **Non-cryptic not identified**: species not identified (possibly identified to genus), with a distinct BIN (or sequence).
5. **Non-cryptic not sequenced identified**: species identified, but not sequenced.
6. **Non-cryptic not sequenced not identified**: species not identified and not sequenced.

Note that the category “cryptic not sequenced” does not exist because species need to have a valid sequence or BIN to demonstrate the status of cryptic species. Here we will discuss categories (1) and (2) and their proportion to the total number of species within each insect taxa studied. Cryptic species may be paired or may include a complex of species which all have ≥2% of similarity among their barcodes (Hebert et al., 2003). One observer (YB) assigned species from all focal groups to the above categories, considering external morphology and molecular data. Study of genitalia was not included because it was not systematically available for all groups and species. The observer was a trained entomologist, has co-authored taxonomic publications, but cannot be considered as an expert in each of the groups studied. However, most of the material considered here has been examined by experts (Table 1). Only adult insects were assigned and, for social insects, only the most relevant caste for taxonomy was considered. For termites, this included the soldier caste (we did not consider the soldierless Apicoterminini, which represents 20 species on BCI; Y. Basset et al., unpubl. data) and for ants, minor workers (when major workers were present). Although 429 species of ants are present on BCI (D. Donoso et al., unpublished data) some are known only by alates collected in Malaise or light traps. To reduce the apparent subjectivity of the method, we include a complete list of cryptic species as recorded in this study, as well as the mean distance divergence between species pairs or among species complexes (Appendix S1).

### Statistical methods

As indicated in the results, some insect assemblages lacked cryptic species. This does not mean that no cryptic species exist in these groups, just that with the local data and sample size in hand, we could not evidence cryptic species in these groups. Hence, some of our statistical analyses considered both datasets including and excluding insect assemblages without cryptic species. To evaluate differences in the proportion of cryptic species among insect assemblages (our first question in the introduction), we considered the range of percentages recorded (categories 1 and 2, see above) among insect assemblages and then performed an ANOVA grouping insect assemblages by orders and trophic guilds (Table 1). Prior the analyses, we did a logit transformation of the results (y=ln(p/(100-p)); groups without cryptic species assigned with p=0). We performed ANOVAs with and without insect assemblages lacking cryptic species.

Regarding our second question, to evaluate phylogenetic distances between insect assemblages, we first considered a comprehensive insect phylogeny, which was constructed hierarchically using c. 440 transcriptomes, 1,490 mitogenomes, and DNA barcodes for 69,000 species (Chesters, 2020). We then selected one representative species from this tree (preferably occurring on BCI or in Panama) for each insect assemblage and pruned the Chesters’ (2020) tree accordingly. Equivalent species for building the phylogeny of insect assemblages are indicated in Table S2 and the final phylogeny of assemblages is displayed in Fig. S1.

We then performed two different analyses, in both cases accounting for phylogeny. First, we modeled the presence of cryptic species (dependent variable: 0 – absent, 1-present) in all insect assemblages (n=22) with a phylogenetic logistic regression for binary dependent variables (Ives & Garland, 2010). Second, we modeled the percentage of cryptic species in insect assemblages with cryptic species (see results; n=11) with a phylogenetic linear regression model (Ho & Ane, 2014). Independent variables for these models are all detailed in Appendix S2 and included the following. *Independent variables related to recent divergence.* The historical age of each taxon was inferred from the age of the earliest known and reliable fossil record, searched in the available literature (Table S2). This was preferred to considering estimates of the ages of diversification of lineages using Bayesian relaxed clock methods (e.g., Wahlberg, 2006), because these methods were not all available for the taxa studied and/or varied a lot in their application. Ages were reported according to the Paleobiology Database (http://paleobiodb.org/) and when reported as ranges, their mid-range was included in models. *Independent variables related to niche conservatism.* Genome size varies tremendously among insect groups and is known to be related to insect growth, metabolism, life history traits and body size (Alfsnes et al., 2017). Estimates of genome size for each assemblage were extracted from Cong et al. (2022). *Independent variables related to morphological convergence*. These included: trophic guild (herbivore, predator, saproxylophagous, pollinator and varia [ants]); main larval habitat (vegetation, wood, soil, nest [social insects]). Dispersal ability was inferred from general ecology and Johnson (1966; low, medium, high). Average body length represents a proxy for dispersal abilities and for each taxon was compiled by considering the mid-range of minimum and maximum values reported by Rainford et al. (2016), unless indicated (Appendix S2). Average adult life span was extracted from Carey (2001: days, weeks, months, year [worker life span for social insects]). *Independent variables related to species pool and sample size*. As an estimate of species pool, we considered regional and local (BCI) species richness (Appendix S2). As variables accounting for sample size, we considered the number of specimens sequenced and its ratio to the overall number of specimens collected. A coarse estimate of local rarity was computed by considering the average percentage occupancy of species within samples for each taxon. Monitoring data on BCI were used for this purpose and reflect whether species are on average well distributed within samples. Taxonomic knowledge may also potentially influence the discovery of cryptic species, as characters to delineate species may be better recognized in well-known taxa. To account for this variable, we first searched Google scholar with search terms “taxonomy” and the insect taxa and recorded the number of references (search performed on 30 March 2023). Second, we considered the ratio between this number of references and regional species richness. Eventually, we also included the % of species identified, as this can also reflect taxonomic knowledge.

The relationships between all dependent variables were explored using a matrix of Pearson correlations (categorical variables were converted to dummy variables to enable this comparison). We removed variables if they were highly correlated (r > 0.7) with other variables. Variables removed included all categorical variables (Guilds, Larval habitat, Dispersal and Lifespan) plus regional species richness. Final selection of variables and their correlation are detailed in Fig. S2. Phylogenetic logistic regressions were performed with the r package “phylolm” (Ho & Ane, 2014). We first modeled a simple logistic binomial regression without accounting for phylogeny with the function *glm* (family=binomial), then used the stepwise function *phylostep* to inform about the best possible inclusion of variables in the model and the function *phyloglm* (with method “logistic_MPLE”) to calculate the final model accounting for phylogeny. The phylogenetic linear regression was performed with the function *phylolm* of the “phylolm” r package, with the Ornstein-Uhlenbeck model with an ancestral state to be estimated at the root. Tests of phylogenetic signals were evaluated with the r package “phylosignal” (Keck et al., 2016). The function *phyloSignal* was used to test for significant phylogenetic signal in the percentage of cryptic species and in the three best variables included in the model (see results), calculating the indices Cmean, I, K, K.star and Lambda (Keck et al., 2016). Local Indicators of Phylogenetic Association (LIPA), based on Moran’s I, were calculated and visualized with the function *phylo4d*, *lipaMoran* and *barplot.phylo4d*, for the percentage of cryptic species and for the three variables included in the model.

Question 3 is problematic because cryptic species were infrequently recorded in samples (see results). Consequently, it was difficult deriving sound variables related to their spatial and temporal distribution. Further, a similar approach than for Question 2 was not possible at the level of cryptic species because (a) no phylogenetic tree was available for the +2,000 BCI species surveyed (see results); and (b) few relevant ecological information was available for cryptic species, which often lacked binomials. Instead, we attempted to document some of the properties of cryptic species available in the context of this study. We first calculated the mean distance divergence between pairs or groups of cryptic species (Appendix S1). This reports the sequence divergence between barcode sequences at (mostly) genus level. It was calculated with the “Distance Summary” analytical tool provided by the Barcode of Life Data System, with parameter distance model = Kimura 2 Parameter, pairwise deletion method and BOLD aligner (Amino Acid based HMM). Kruskal-Wallis and Dunn tests were used to test differences between insect orders, assemblages and guilds. Second, we summarized the information available on the relative abundance of cryptic species and the feeding guilds to which they belong. For butterflies, which arguably represent the better-known taxon, we also tested for differences in the host specificity, geographic range and forewing length between cryptic and non-cryptic species (Appendix S3) with t-tests. Butterfly data were extracted from Basset et al. (2015).

To estimate the number of cryptic species present in our study system (Question 4), we calculated asymptotic estimates with the R package iNext (Hsieh et al., 2020), by considering the rate of accumulation of cryptic vs. non-cryptic species in all insect assemblages in function of the number of individuals sequenced. We extrapolated data to about twice the number of individuals sequenced (ca. 20,000). We compared the percentage of cryptic species to the total number of species sequenced either with the observed data or with the asymptotic estimator.

## RESULTS

A total of 326,412 individuals representing 2,006 local species occurring on BCI are considered in this contribution. Although only 7,544 individuals were sequenced (2.3% of total individuals), this covered 76.1% of all species, out of which 56.9% were identified to species level (Tables 1 and S2). We observed 96 cases of cryptic species, representing in total 214 species, 78 cases involving species pairs, 14 involving triplets and 4 involving quadruplets (Appendix S1; illustrated examples in Appendix S4). Overall, the proportion of cryptic species was 10.6% when considering all species scored (Fig. 1; exact values in Table S3). Mean percentage per insect assemblage was 6.0±1.5% (s.e., n=22) and 12.1±1.3% (n=11) of cryptic species when removing insect assemblages without cryptic species. Percentages of cryptic species per taxa ranged from 0 to 19% (Fig. 1; Table S3). Half of assemblages studied lacked cryptic species and high percentages approaching 20% of cryptic species were observed in Pieridae and Formicidae. Genera with a relatively high number of cryptic species recruited from Formicidae (8 genera with 3 or more cryptic species; especially *Solenopsis* and *Pheidole*; Appendix S3), Crambidae (5 genera including *Diaphana* and *Omiodes*), Geometridae (5 genera including *Cyclophora* and *Pleuroprucha*), Hesperiidae (3 genera including *Panoquina*, *Calpodes* and *Astraptes*) and Pyralidae (2 genera including *Paramacna* and *Deuterollyta*).

**FIGURE 1.**
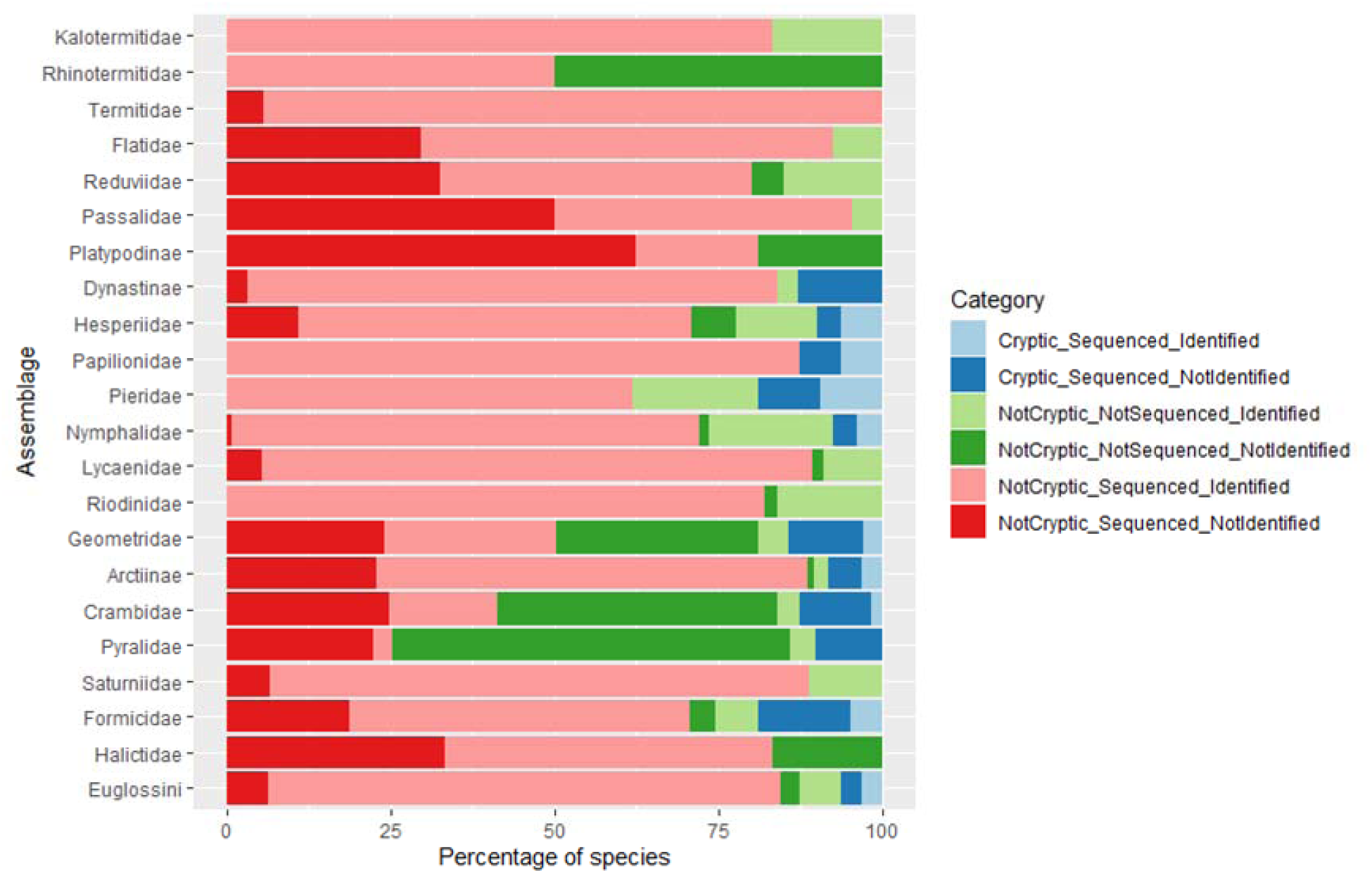
Percentage of the number of species recorded in each category, detailed for each assemblage studied.

None of the ANOVAs that we performed when grouping insect assemblages by orders or guilds were significant (all with p >0.05; the closest test to significance was comparing insect assemblages by order, accounting for assemblages without cryptic species, F_4,17_=2.3, p=0.108).

Regarding Question 2, the percentage of cryptic species within insect taxa was not or weakly influenced by phylogeny (Table S4). The stepwise phylogenetic logistic regression indicated that the best variables in the model were local species richness, ratio of knowledge to regional richness (both with a positive coefficient) and historical age (negative coefficient). Our final model with the lowest AIC included only local species richness, ratio of knowledge to regional richness and explained 62% of variation in the occurrence of cryptic species within insect assemblages (Table 2). The plot of these two independent variables in relation to the occurrence of cryptic species within insect assemblages is illustrated in Fig. S3. LIPA indicators confirmed that significant phylogenetic signals for each tip of the tree were rare in variables percentage of cryptic species, local species richness and ratio of knowledge to regional richness, despite being frequent in historical age (Fig. S4). However, the best phylogenetic linear model accounting for the percentage of cryptic species within insect assemblages with cryptic species was not significant, with historical age as the sole variable approaching significance (Table 2).

**TABLE 2.** Best models for (a-c) logistic regressions with the occurrence of cryptic species within insect assemblages as dependent variable; (d) phylogenetic regression with the percentage of cryptic species within insect assemblages as dependent variable. alpha = phylogenetic correlation parameter for the phylogenetic logistic regression.

| Model/Variable | AIC /<br>Coefficient | R2 / s.e. | alpha / z | p |
| --- | --- | --- | --- | --- |
| *(a) Logistic binomial regression* | 22.4 | 0.462 | - | - |
| Intercept | -3.6288 | 1.5825 | -2.2930 | 0.0218 * |
| Local species richness | 0.0393 | 0.0192 | 2.0440 | 0.0409 * |
| Ratio knowledge to regional richness | 0.2442 | 0.1334 | 1.8310 | 0.0671 |
| *(b) Stepwise phylogenetic logistic regression* | 29.5 | 0.483 | - | - |
| Intercept | 0.3836 | 0.3625 | 1.0582 | 0.3040 |
| Local species richness | 0.0029 | 0.0008 | 3.5569 | 0.0023 ** |
| Ratio knowledge to regional richness | 0.0408 | 0.0172 | 2.3731 | 0.0289 * |
| Age | -0.0062 | 0.0032 | -1.9607 | 0.0656 |
| *(c) Best phylogenetic logistic regression* | 24.7 | 0.620 | 1.567 | - |
| Intercept | -3.6272 | 1.6629 | -2.1813 | 0.0292 * |
| Local species richness | 0.0241 | 0.0107 | 2.2543 | 0.0242 * |
| Ratio knowledge to regional richness | 0.2188 | 0.1230 | 1.7787 | 0.0753 |
| *(d) Best phylogenetic regression* | 65.9 | 0.291 | - | - |
| Intercept | 6.7350 | 2.9904 | 2.2522 | 0.0508 |
| Age | 0.1121 | 0.0583 | 1.9230 | 0.0866 |

Concerning Question 3, the mean distance divergence between pairs or groups of cryptic species was 8.97 % ± 0.63 (range 1.07-33.13 %, n=96 cases; Appendix S1). For the insect orders, families and guilds for which we had sufficient data (number of cryptic species), there were significant differences between mean distance divergence in taxa and guilds (Fig. 2). However, these differences were driven mostly by ants, which showed rather high mean distance divergence as opposed to other insect families (Fig. 2). Without surprise, mean distance divergence was significantly higher when comparing cases involving species pairs to cases involving triplets and quadruplets (Mann-Whitney test, U=404.5, p <0.01; mean pairs = 8.03 % ± 0.61; mean triplets and quadruplets = 13.04 % ± 1.82).

**FIGURE 2.**
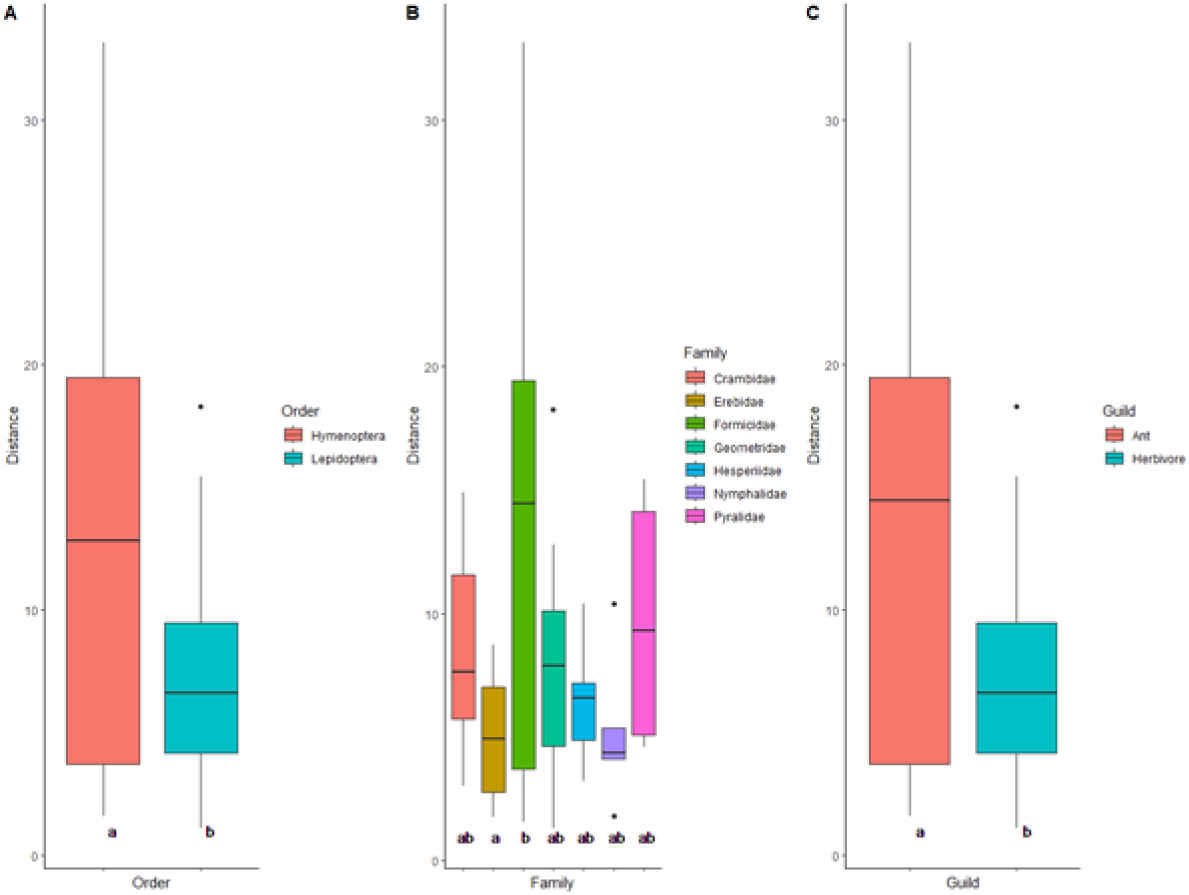
Mean distance divergence detailed for pairs or groups of cryptic species belonging to A) orders, B) families and C) guilds, when sufficient sample size allows comparisons. Different letters for groups denote significant differences (p<0.05) after Kruskal-Wallis or Dunn tests.

All pairs of cryptic species (or species complexes) were congenerics (Appendix S3). The percentage of cryptic species present in each insect guild increased along the series predators (0%) – saproxylophagous (4.1%) – pollinators (5.3%) – herbivores (9.9%) – ants (18.8%; Appendix S3). The abundance of cryptic species was low in our collections: cryptic individuals sequenced represented 14.2% of the total number of individuals sequenced and only 0.3% of the total number of individuals collected (Table 1, Appendix S3). Neither the index of host specificity, nor the index of geographic range, nor the forewing length of butterflies differed significantly between cryptic and non-cryptic species (t-tests, all with p >0.50).

Eventually, relating to Question 4, the rate of species accumulation for non-cryptic species sequenced was much steeper than for cryptic species (Fig. 3). Asymptotic estimates were 291.5 ± 26.9 (s.e.) for cryptic species and 2,272.0 ± 63.6 for non-cryptic species. This asymptotic total of ca. 2,564 species (to compare with 2,006 species observed, Table 2) was reached after about 40,000 individuals sequenced, about four time the number of individuals actually sequenced. With asymptotic estimates, the percentage of cryptic species to total number of species was 11.4% (compare with 10.6% for cryptic species observed, Table S3).

**FIGURE 3.**
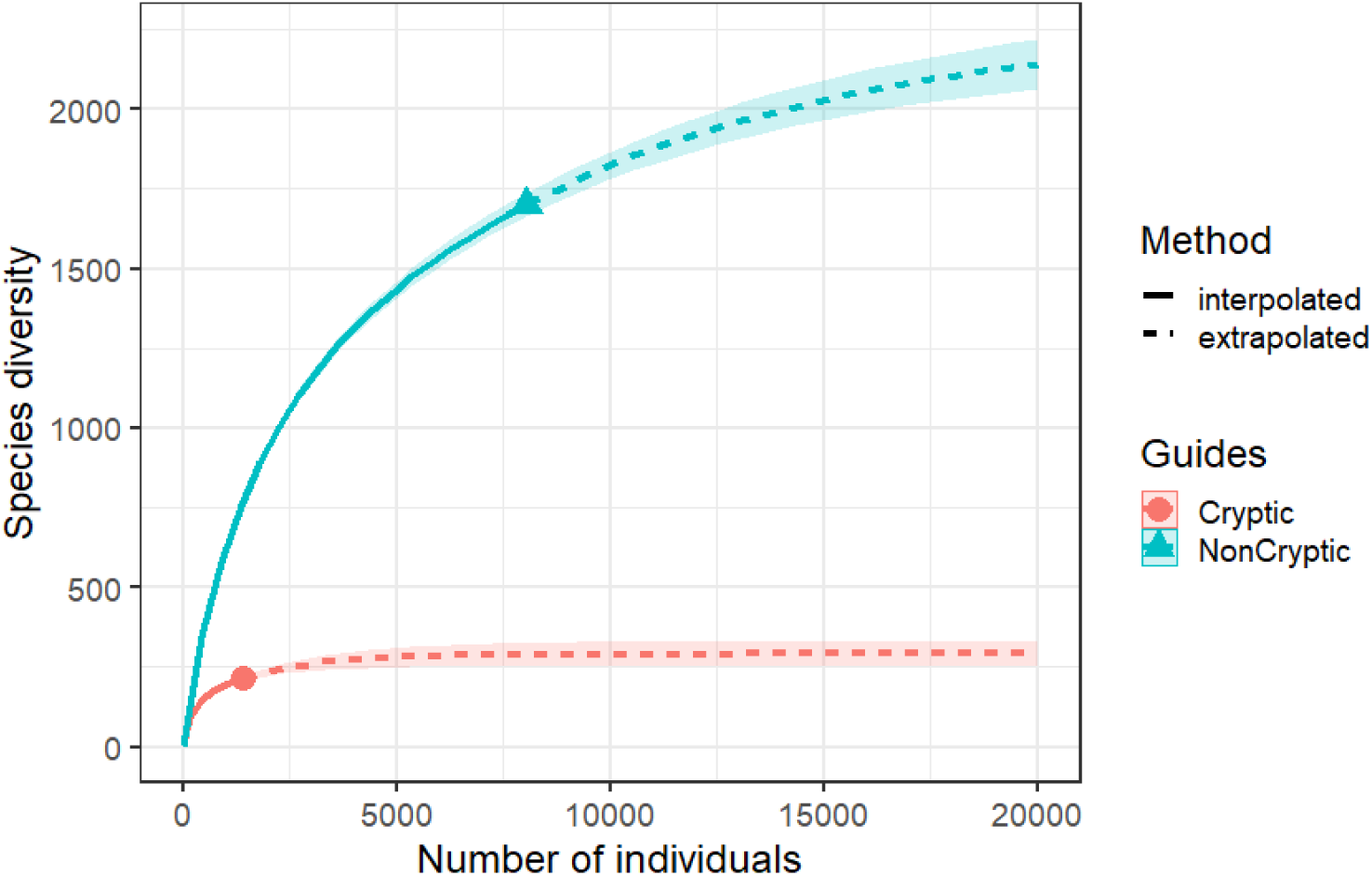
Accumulation of species richness for cryptic species and non-cryptic species as function of the number of individuals sequenced.

## DISCUSSION

As far as we are aware, this is the first contribution discussing local cryptic diversity for a wide range of insect assemblages at a tropical location (Table S1). After scoring 2,006 species including predators, saproxylophagous, pollinators, herbivores and ants recruiting from 22 different assemblages originating from BCI in Panama, we detected 214 cryptic species (10.6% of the local pool of species studied). These cryptic species were all sequenced and had a distinct barcode index number but were indistinguishable from another species when considering external morphology. The % of cryptic species varied greatly among assemblages (0-19%), with half of assemblages devoid of cryptic species, and the highest proportions of cryptics in Pieridae and Formicidae. The % of cryptic species among assemblages was weakly influenced by phylogeny and it was best explained by the local number of species and an index of taxonomic knowledge (62% of variance explained). However, the model explaining the % of cryptic species in assemblages with cryptic species was not significant. The characteristics of cryptic species, which overall were distinguished by an average distance divergence in sequence of 9%, remain unclear. Asymptotic estimates indicated that cryptic species may represent 11.4% of species on BCI, for the assemblages studied.

Our study was limited to the focal taxa monitored by the ForestGEO Arthropod Initiative on BCI. These taxa were in part selected for tractable taxonomy, hence other insect taxa with lower taxonomic knowledge may well prove to include higher shares of cryptic species. On the other hand, individuals sequenced may have been biased positively towards cryptic species since, to improve our reference collections, we attempted to sequence in priority potential complexes of species. Only external morphology was considered to score cryptic species and some of these species may be separated by the examen of genitalia or ecological information (host plants, bioacoustics, etc.). We also note that morphological characteristics, such as color for example, may be better evidenced in live specimens as opposed to dead specimens in collection. Our scoring system may appear subjective, but we provide a full list of the cryptic species that may be critically revised, and we also used mean distance divergence in the sequences of species to characterize pairs or complexes of cryptic species.

In reference to Question 1, our estimate of 10.6% of cryptic species for the assemblages studied on BCI is certainly higher than that proposed by Stork (1-2%), but also much lower than that suggested by Li and Wiens (2023: 310%). Our local study may include a low percentage of cryptic species, especially when comparing regional studies (Table S1), but is useful to compare the incidence of cryptic species among insect assemblages. The proportion of cryptic species in insects appears to be scale-dependent, as regional studies may include different habitats likely to promote reproductive isolation and allopatric speciation. For example, we report 0% of cryptic species on BCI for Rhinotermitidae, Termitidae, Reduviidae, Lycaenidae and Saturniidae, but we know of verified (i.e., with a barcode index number) cryptic species in all these groups in the whole country of Panama (Basset et al., unpubl. data). Also compare our results for Saturniidae with Janzen et al. (2012), who reported 34.7% cryptic species for this group in a much larger geographic area encompassing different habitats (dry, lowland and montane forest) than the 1,500ha BCI. Further, the results of our study do not support the argument of Pfenninger & Schwenk (2007) that cryptic species are evenly distributed among taxa. In this context, we may ask which insect taxa on BCI, not surveyed in this study, may support a high share of cryptic species? Since our logistic regressions suggested that the richness of the local species pool was positively correlated with the percentage of cryptic species within assemblages, we expect a high number of cryptic species in species-rich families, such as Curculionidae, Staphylinidae, Noctuidae, Cecidomyiidae, Braconidae, etc. (e.g., Srivathsan et al. 2023). Taxa thriving in more extreme environments may also support a high share of cryptic species (Bickford et al., 2007) and on BCI, this may be relevant to specialized herbivores and parasitoids, especially in the forest canopy, where, for example, insects may have to experience desiccation and stressful temperatures (Bujan *et al*., 2016).

Relevant to Question 2, among the hypotheses that Mayew (2007) reviewed to account for the high species richness in insects, six may be particularly pertinent to cryptic species: historical age, speciation rates, extinction rates, ecological specialization, sexual selection, and genetic versatility. Further, recent divergence, niche conservatism and morphological convergence, may also represent important concepts to explain the incidence of cryptic species (Fišer et al., 2018). Unfortunately, most of the variables related to these premises are rarely available for insect species, even less so for tropical species. This issue is even more challenging when attempting to derive representative and quantitative variables at the level of higher insect taxa, such as in this contribution. Our variables accounting for niche conservatism or morphological convergence (Appendix S2), were often crude, represented as categorical variables which proved to be of limited value in our analyses (larval habitat, dispersal ability, lifespan), or simply unavailable (gene flow, molecular evolutionary rates, etc.). Nevertheless, we could account for the effect of phylogeny on the incidence of cryptic species in insect assemblages. It appeared to be weak, as compared to, for example, its effect on the variable historical age (Fig. S4). Significant phylogenetic signals influencing the incidence of cryptic species were only detected for Geometridae and Papilionidae (LIPA, Fig. S4). This reflects the concentration of genera with cryptic species in these two families (Appendix S3), not their relative proportion of cryptic species.

The richness of the local species pool represented the best variable explaining the incidence of cryptic species within assemblages. This positive relationship was expected as a surrogate of the sum of several factors (cf. above) favoring locally high species richness, although the proximate factor accounting for the incidence of cryptic species remains unclear. Since tropical insect assemblages are often more diverse than temperate ones (e.g., Basset et al., 2015), this confirms that the former may also support a higher share of cryptic species than the later (Table S1). Of interest, one of the variables accounting for taxonomic knowledge (ratio of taxonomic knowledge to regional richness) was also important in our logistic regressions. This implies that a better taxonomic knowledge (relative to the number of species described) may result in a better knowledge of species limits and of their morphological characteristics, and, ultimately, a better resolution in the delineation of cryptic species. The (weak) negative influence of historical age in our logistic regressions was expected, as recent taxa divergence often favors cryptic diversity (Bickford et al., 2007; Fišer et al., 2018; Struck et al., 2018).

However, the lack of obvious influence of variables accounting directly for sampling effort, such as the number of specimens sequenced, was unexpected. As noted previously, our selection of specimens to be sequenced, which may have been biased towards potential cryptic species, may explain this situation. We note that, at current pricing (Canadian Centre for DNA Barcoding, pers. comm.), sequencing each specimen collected in this study would have cost about $3.5 mio. A more informed selection of potential cryptic species including about 5,000 individuals would have probably detected most cryptic species present within the insect assemblages studied (see Fig. 3), at current pricing of ca. $52K.

Related to Question 3, it was challenging to characterize cryptic species with the data in hand. We observed that most cryptics are congeneric species with an average mean distance divergence of 9% between them. Unexpectedly, mean distance divergence between cryptic species was high within Formicidae, a family with a high percentage of cryptic species. Since divergence was calculated within genera, this suggests that some of the ant genera with many cryptic species (e.g., *Pheidole*, *Solenopsis*, etc.; Appendix S3) may be poorly delimited and in need in taxonomic revision. Looking at the incidence of cryptic species within all individuals sequenced, our data suggest that population levels of cryptic species may be low relative to that of non-cryptic species. This should be confirmed by more exhaustive studies. Beside the variables that we already mentioned to explain the incidence of cryptic species within assemblages, additional variables may be more relevant at the species level but could not be investigated in this contribution. For example: (a) rapid barriers to reproductive isolation, such as the infestation rate of endosymbiotic intracellular *Wolbachia* spp. bacteria (Zhang et al., 2022), sexual selection (Jones, 1997), or allopatry (Lukhtanov et al., 2015); (b) niche separation on an axis that does not include morphological factors (bioacoustics, host plants, etc., Bickford et al., 2007); (c) molecular issues such as mitochondrial-nuclear discordance in molecular data or mtDNA sweeps (Belfiore *et al*., 2013; Hinojosa et al., 2019; Li et al., 2021); or (d) porous species boundaries with extensive hybridization (D’Ercole et al., 2021). All these issues may result in problematic resolution of species, potential misidentifications, and high apparent incidence of cryptic species. Examples abound in entomological literature, particularly with the better studied butterflies. For example, see the community study of D’Ercole et al. (2021) and the discussion about the pierid genus *Phoebis* in Nunez et al. (2021), which includes two cryptic species on BCI.

With respect to Question 4, our estimates of 10.6% of cryptic species for the local pool of species studied, with further an asymptotic estimator suggesting 11.4% of cryptic species are worth discussing for the whole of BCI. There are several arguments to push these estimates up or down. In support of higher estimates, first we note that the number of individuals which were assigned to cryptic species (and were sequenced by definition) represented 14.2% of all individuals sequenced. Second, not all species of assemblages studied were collected on BCI and not all individuals were sequenced. Our regressions suggest that the richness of the local species pool is important to explain the incidence of cryptic species, less so sampling effort per se (number of individuals sequenced). Many of the species-rich insect families occurring on BCI were not evaluated in our study, so the upper bound of the percentage of cryptic species is difficult to estimate. Conversely, estimates close to what we observed, or lower estimates of cryptic species are also plausible. We noted that the % of cryptic species overall was relatively independent of phylogeny, so this may represent an argument to valid our observations for the assemblages studied to the whole of the insect community of BCI. Since small effective population size and occasional nuclear insertion of mtDNA can lead to multiple copies of COI, estimates of cryptic species may also be pushed down when cryptic species are detected solely by mitochondrial molecular data as opposed to nuclear data (Li et al., 2021). However, in this study, sources of genetic variability should be devoid of geographic variability and, in particular, mitochondrial introgression should be reduced (Bastos□Silveira *et al*., 2012). We also noted (Appendix S1) that different BINs were sometime,es (but rarely) defined with < 2% of sequence similarity, a usual threshold used to delineate species (Hebert *et al*., 2003). Another argument, particularly for some ant species, is that some genera lack a modern taxonomic revision that could be used by relatively “untrained” observers. All these arguments could push down our estimates. However, on balance, we consider that our observed estimates of cryptic species probably represent lower bounds to the true cryptic diversity on BCI, simply because of the highly speciose taxa that were not investigated in this study.

If anything, this study provided hints as to which species complexes may be worth further taxonomic and molecular studies. However, as indicated by this study, an additional ten to fifteen percent (and possibly more) of cryptic species to a local pool of insect species in the tropics represents a significant contribution to biodiversity, which should not be ignored, particularly in conservation planning. It would be stimulating to obtain similar estimates of cryptic species for temperate forests, as they may prove to be lower than in this study. Unfortunately, as this contribution confirms, cryptic insect species are difficult to study in the tropics, for lack of ecological information and given the relatively low interest in insect as opposed to vertebrate conservation (Leather, 2009). Our data also suggest that the insect populations of cryptic species may be rather low relative to non-cryptic species. If this is confirmed by more exhaustive studies, insect populations of cryptic species in the tropics may be particularly vulnerable to land use and climate changes (for a similar argument for aquatic temperate insects, see Bálint *et al*., 2011).

## AUTHOR CONTRIBUTIONS

**Yves Basset:** conceptualization; investigation; data curation; writing. **Greg P. A. Lamarre, David A. Donoso, Daniel Souto-Vilarós, Héctor Barrios:** investigation; writing. **Filonila Perez, Ricardo Bobadilla, Yacksecari Lopez, José Alejandro Ramírez Silva**: data curation.

## ACKNOWLEDGMENTS

We thank the Smithsonian Tropical Research Institute for logistical support and all experts cited in Table 1 for taxonomic help.

## DATA AVAILABILITY STATEMENT

Data are available as Supplementary file Appendix S1.

## SUPPORTING INFORMATION

**APPENDIX S1** List of cryptic species recorded on BCI for the insect assemblages studied (Excel file, tab 1).

**APPENDIX S2** List of independent variables used in models explaining the occurrence of cryptic species, detailed for each insect assemblages studied (Excel file, tab 2).

**APPENDIX S3** List of traits available for each cryptic species recorded in this study (Excel file, tab 3).

**APPENDIX S4.**
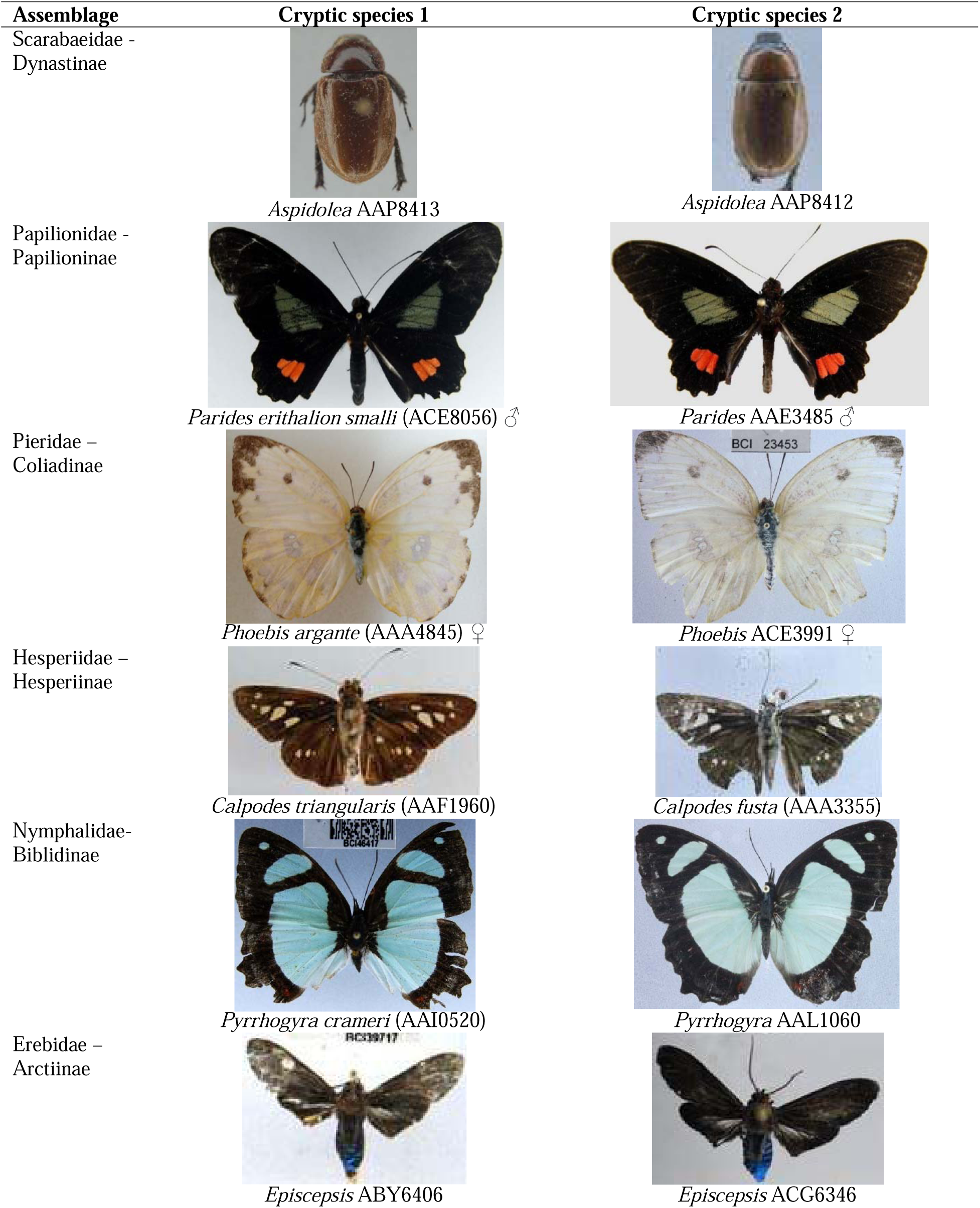
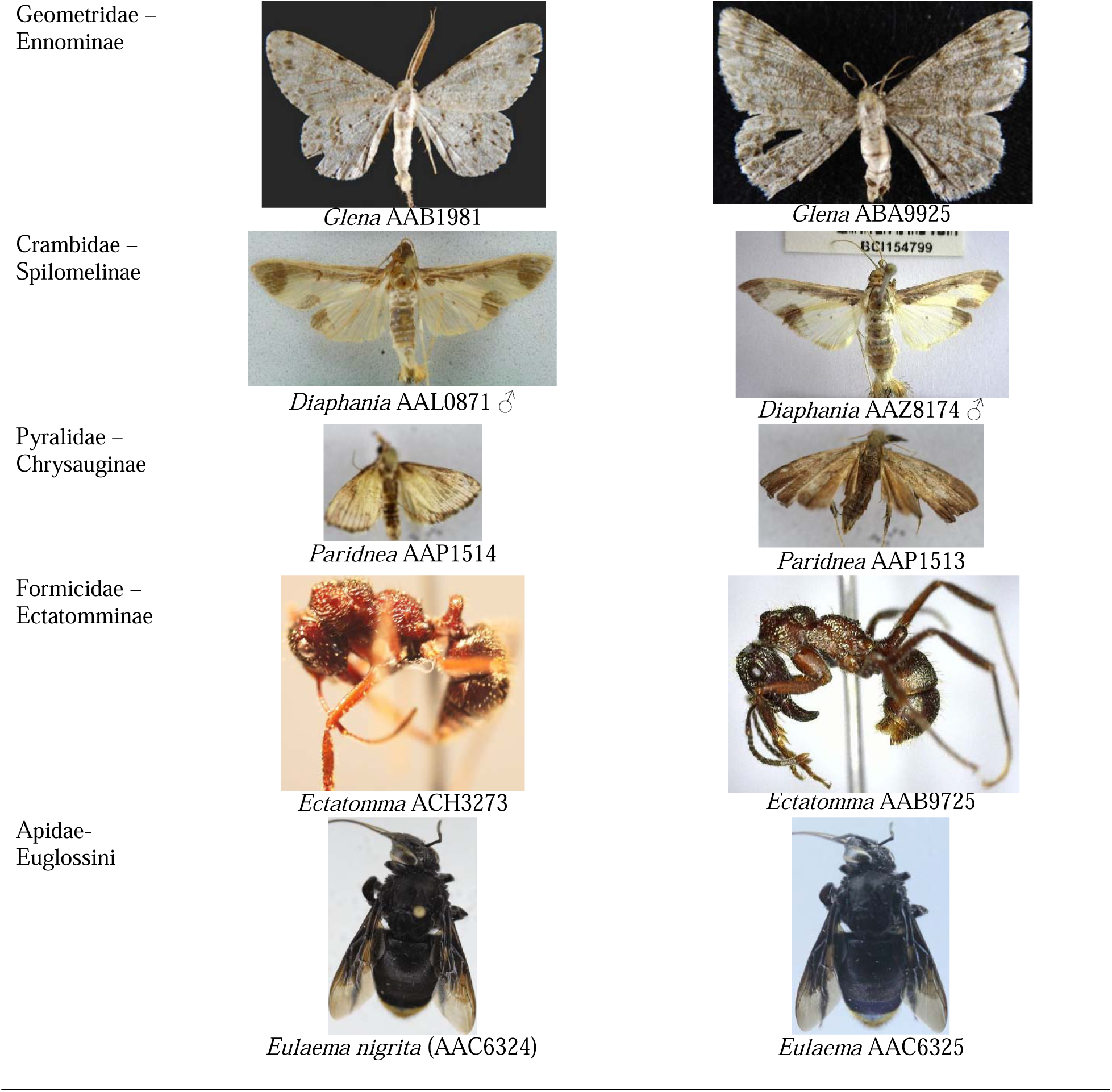
Examples of pairs of cryptic species (habitus) for each insect assemblage studied.

**TABLE S1.**
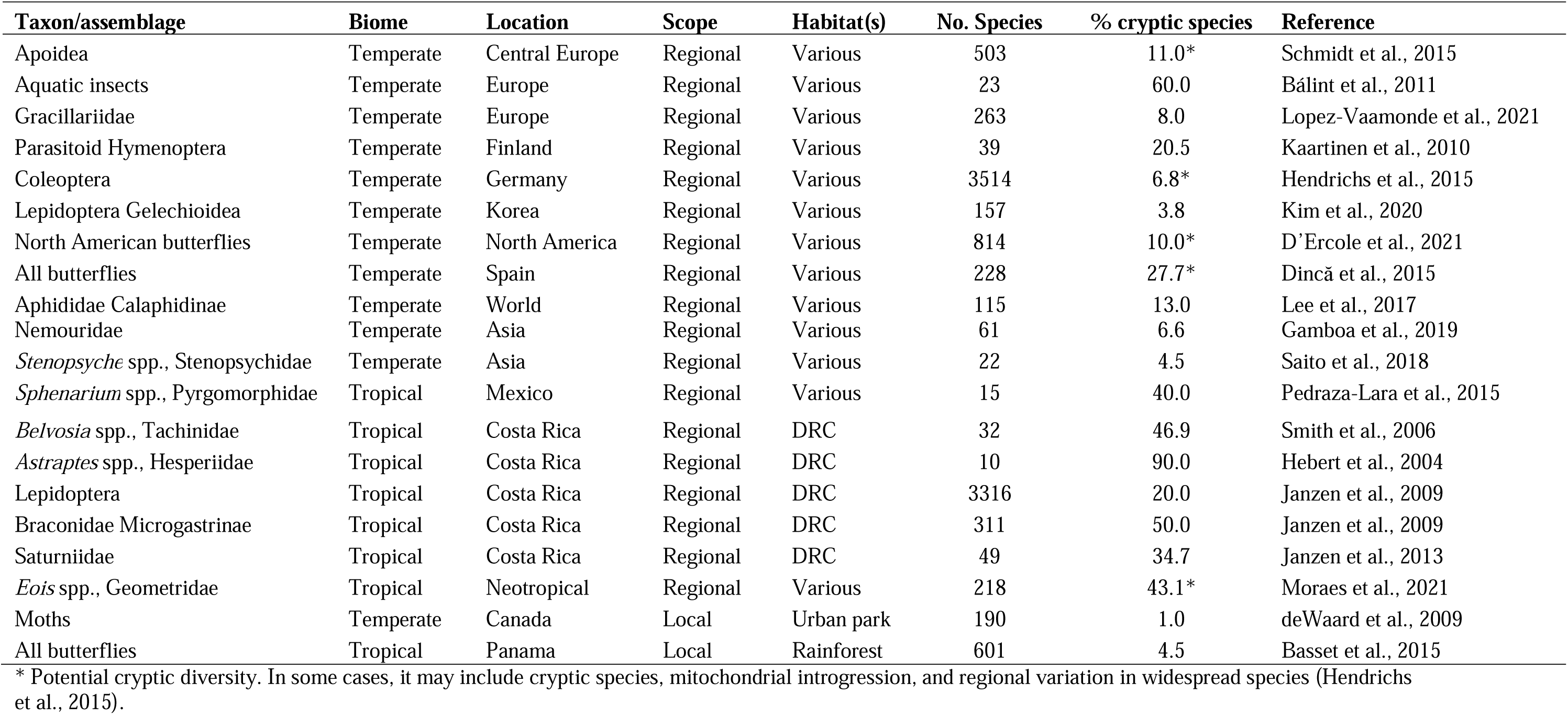
Percentage of cryptic species reported in selected insect studies performed at the community level (including at least 10 species). The number of species refers to the total number of species (including cryptic species) covered by the study. DRC = Dry, rain and cloud forests.

---

**TABLE S2.** Species from Chesters (2020) used as equivalent for building the phylogeny of focal assemblages.

| **Assemblage** | **Family** | **Subfamily** | **Equivalent species** | **Equivalent family** | **Equivalent subfamily** |
| --- | --- | --- | --- | --- | --- |
| Kalotermitidae | Kalotermitidae | *** | *Calcaritermes temnocephalus* | Kalotermitidae | *** |
| Rhinotermitidae | Rhinotermitidae | *** | *Coptotermes testaceus* | Rhinotermitidae | *** |
| Termitidae | Termitidae | *** | *Microcerotermes arboreus* | Termitidae | Termitinae |
| Flatidae | Flatidae | *** | *Flatormenis proxima* | Flatidae | *** |
| Reduviidae | Reduviidae | *** | *Brontostoma colossus* | Reduviidae | *** |
| Passalidae | Passalidae | *** | *Lucanus cervus* | Lucanidae | *** |
| Platypodinae | Curculionidae | Platypodinae | *Platypus calamus* | Curculionidae | Platypodinae |
| Dynastinae | Scarabaeidae | Dynastinae | *Cyclocephala melanocephala* | Scarabaeidae | Dynastinae |
| Hesperiidae | Hesperiidae | *** | *Astraptes alardus* | Hesperiidae | Eudaminae |
| Papilionidae | Papilionidae | *** | *Parides erithalion* | Papilionidae | Papilioninae |
| Pieridae | Pieridae | *** | *Eurema albula* | Pieridae | Coliadinae |
| Nymphalidae | Nymphalidae | *** | *Philaethria dido* | Nymphalidae | Heliconiinae |
| Lycaenidae | Lycaenidae | *** | *Lycaena thetis* | Lycaenidae | Lycaeninae |
| Riodinidae | Riodinidae | *** | *Mesosemia hesperina* | Riodinidae | Riodininae |
| Geometridae | Geometridae | *** | *Abraxas leopardina* | Geometridae | Ennominae |
| Arctiinae | Erebidae | Arctiinae | *Virbia mentiens* | Erebidae | Arctiinae |
| Crambidae | Crambidae | *** | *Parapoynx fluctuosalis* | Crambidae | Acentropinae |
| Pyalidae | Pyalidae | *** | *Deuterollyta basilata* | Pyalidae | Epipaschiinae |
| Saturniidae | Saturniidae | *** | *Saturnia albofasciata* | Saturniidae | Saturniinae |
| Formicidae | Formicidae | *** | *Rogeria belti* | Formicidae | Myrmecinae |
| Halictidae | Halictidae | *** | *Halictus gemmeus* | Halictidae | *** |
| Euglossini | Apidae | Apinae-Euglossini | *Euglossa imperialis* | Apidae | Apinae-Euglossini |

**TABLE S3.** Details of the number of local species recorded in each category and assemblage. In brackets, % of the total number of local species.

| Taxa/Assemblage | CRYPTIC |  | NON-CRYPTIC |  |  |  |
| --- | --- | --- | --- | --- | --- | --- |
|  | *Sequenced* |  | *Sequenced* |  | *Not sequenced* |  |
|  | Identified | Not identified | Identified | Not identified | Identified | Not identified |
| Kalotermitidae | 0 (0) | 0 (0) | 5 (83.3) | 0 (0) | 1 (16.7) | 0 (0) |
| Rhinotermitidae | 0 (0) | 0 (0) | 2 (50) | 0 (0) | 0 (0) | 2 (50) |
| Termitidae (1) | 0 (0) | 0 (0) | 17 (94.4) | 1 (5.6) | 0 (0) | 0 (0) |
| Flatidae | 0 (0) | 0 (0) | 17 (63.0) | 8 (29.6) | 2 (7.4) | 0 (0) |
| Reduviidae | 0 (0) | 0 (0) | 41 (47.7) | 28 (32.5) | 13 (15.1) | 4 (4.7) |
| Passalidae | 0 (0) | 0 (0) | 10 (45.5) | 11 (50.0) | 1 (4.5) | 0 (0) |
| Platypodinae | 0 (0) | 0 (0) | 3 (18.8) | 10 (62.4) | 0 (0) | 3 (18.8) |
| Dynastinae | 0 (0) | 4 (12.9) | 25 (80.7) | 1 (3.2) | 1 (3.2) | 0 (0) |
| Hesperiidae | 11 (6.4) | 6 (3.5) | 102 (59.7) | 19 (11.1) | 21 (12.3) | 12 (7.0) |
| Papilionidae | 1 (6.3) | 1 (6.3) | 14 (87.4) | 0 (0) | 0 (0) | 0 (0) |
| Pieridae | 2 (9.5) | 2 (9.5) | 13 (62.0) | 0 (0) | 4 (19.0) | 0 (0) |
| Nymphalidae | 5 (3.8) | 5 (3.8) | 95 (71.3) | 1 (0.8) | 25 (18.8) | 2 (1.5) |
| Lycaenidae | 0 (0) | 0 (0) | 47 (83.9) | 3 (5.4) | 5 (8.9) | 1 (1.8) |
| Riodinidae | 0 (0) | 0 (0) | 46 (82.1) | 0 (0) | 9 (16.1) | 1 (1.8) |
| Geometridae | 8 (3.0) | 30 (11.4) | 69 (26.1) | 64 (24.2) | 12 (4.5) | 81 (30.8) |
| Arctiinae | 6 (3.1) | 10 (5.2) | 127 (65.8) | 44 (22.8) | 4 (2.1) | 2 (1.0) |
| Crambidae | 6 (1.8) | 37 (10.9) | 56 (16.5) | 84 (24.7) | 11 (3.2) | 146 (42.9) |
| Pyrilidae | 0 (0) | 11 (10.3) | 3 (2.8) | 24 (22.4) | 4 (3.7) | 65 (60.8) |
| Saturniidae | 0 (0) | 0 (0) | 37 (82.2) | 3 (6.7) | 5 (11.1) | 0 (0) |
| Formicidae | 17 (4.8) | 50 (14.0) | 184 (51.8) | 67 (18.8) | 24 (6.7) | 14 (3.9) |
| Halictidae | 0 (0) | 0 (0) | 3 (50.0) | 2 (33.3) | 0 (0) | 1 (16.7) |
| Euglossini | 1 (3.1) | 1 (3.1) | 25 (78.1) | 2 (6.3) | 2 (6.3) | 1 (3.1) |
| All | 57 (2.8) | 157 (7.8) | 941 (46.9) | 372 (18.6) | 144 (7.2) | 335 (16.7) |

(1) Without Apicoterminae

**TABLE S4.** Indices of phylogenetic signal calculated for the percentage of cryptic species within insect assemblages, as well as for the three best variables included in models (p included in parenthesis). Significant values are indicated in bold.

| Variable | Cmean | I | K | K.star | Lambda |
| --- | --- | --- | --- | --- | --- |
| % cryptic species | 0.086 (0.186) | -0.008 (0.069) | 0.646 (0.077) | 0.655 (0.128) | 0.243 (0.396) |
| Local species richness | 0.075 (0.180) | -0.013 (0.075) | 0.551 (0.254) | 0.598 (0.312) | 0.113 (0.733) |
| Ratio knowledge to regional richness | -0.005 (0.328) | -0.037 (0.248) | 0.468 (0.579) | 0.538 ( 0.525) | 0.028 ( 0.960) |
| Age | 0.129 (0.141) | **0.038 (0.010)** | **1.000 ( 0.004)** | **0.889 (0.006)** | **0.650 (0.038)** |

**FIGURE S1.**
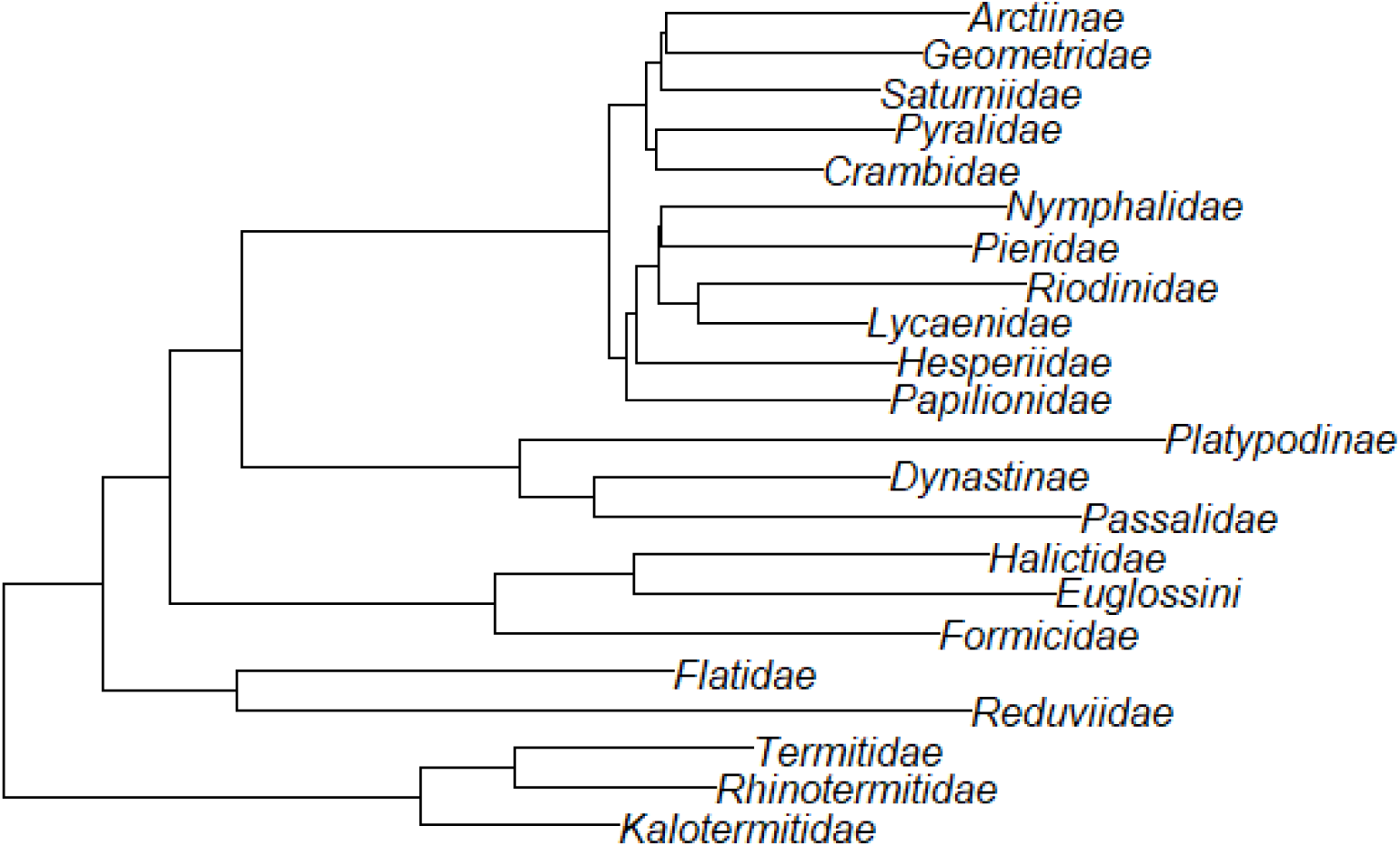
Phylogeny of focal insect assemblages after pruning the tree of Chesters (2020).

**FIGURE S2.**
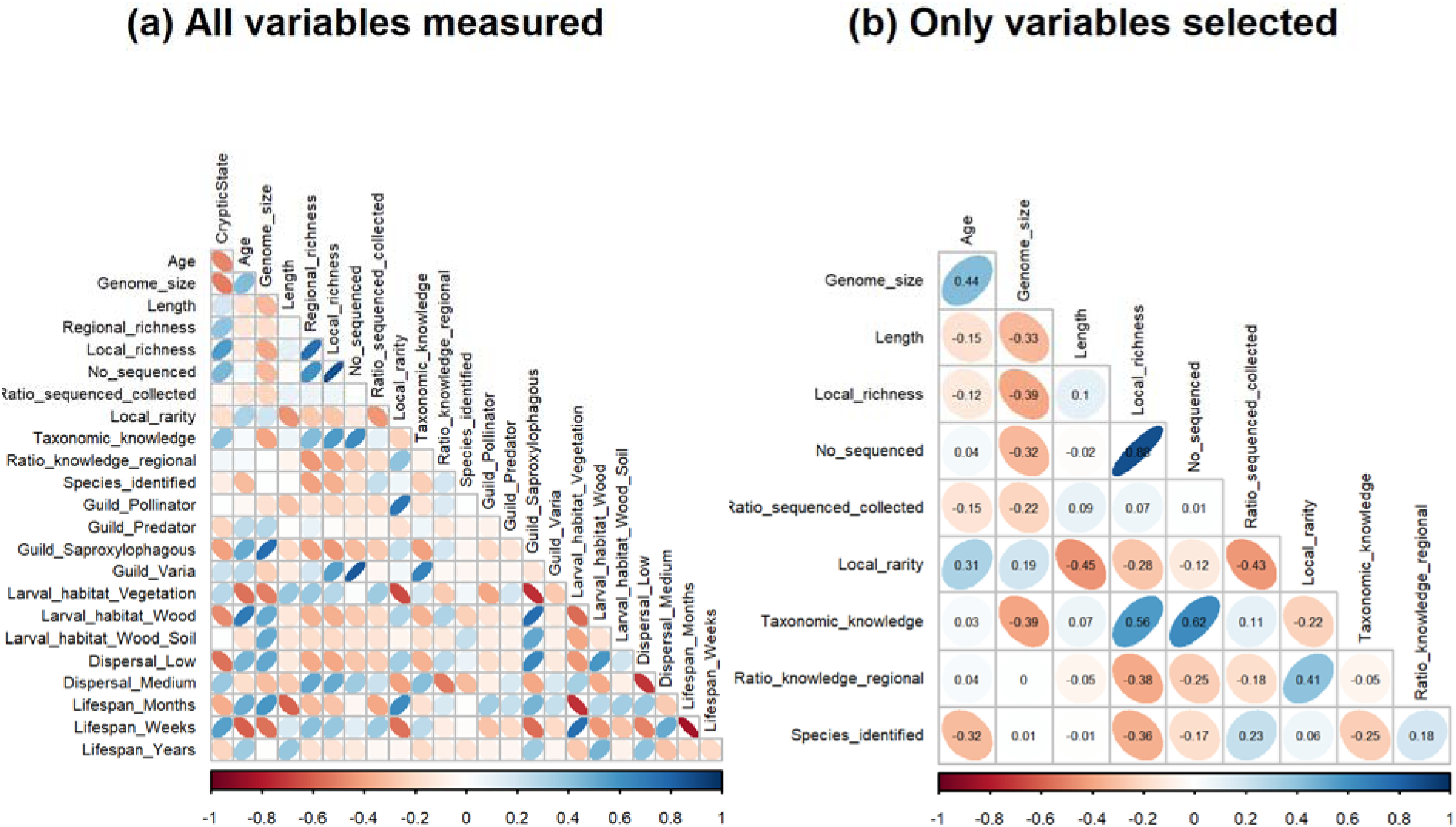
Correlations (Pearson r) between independent variables: (a) all variables measured; (b) only variables selected for analyses.

**FIGURE S3.**
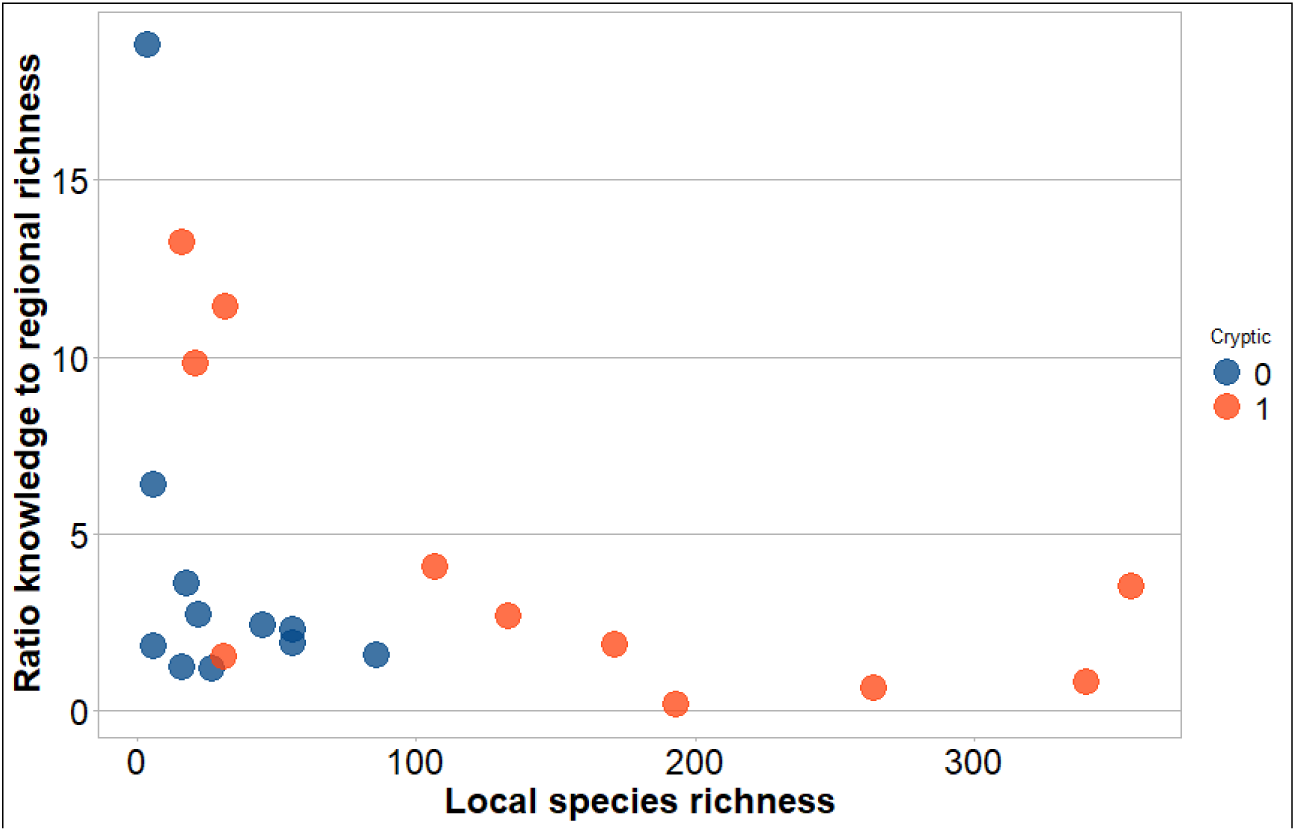
Plot of local species richness vs. the ratio of knowledge to regional richness in relation to the occurrence of cryptic species within insect assemblages.

**FIGURE S4.**
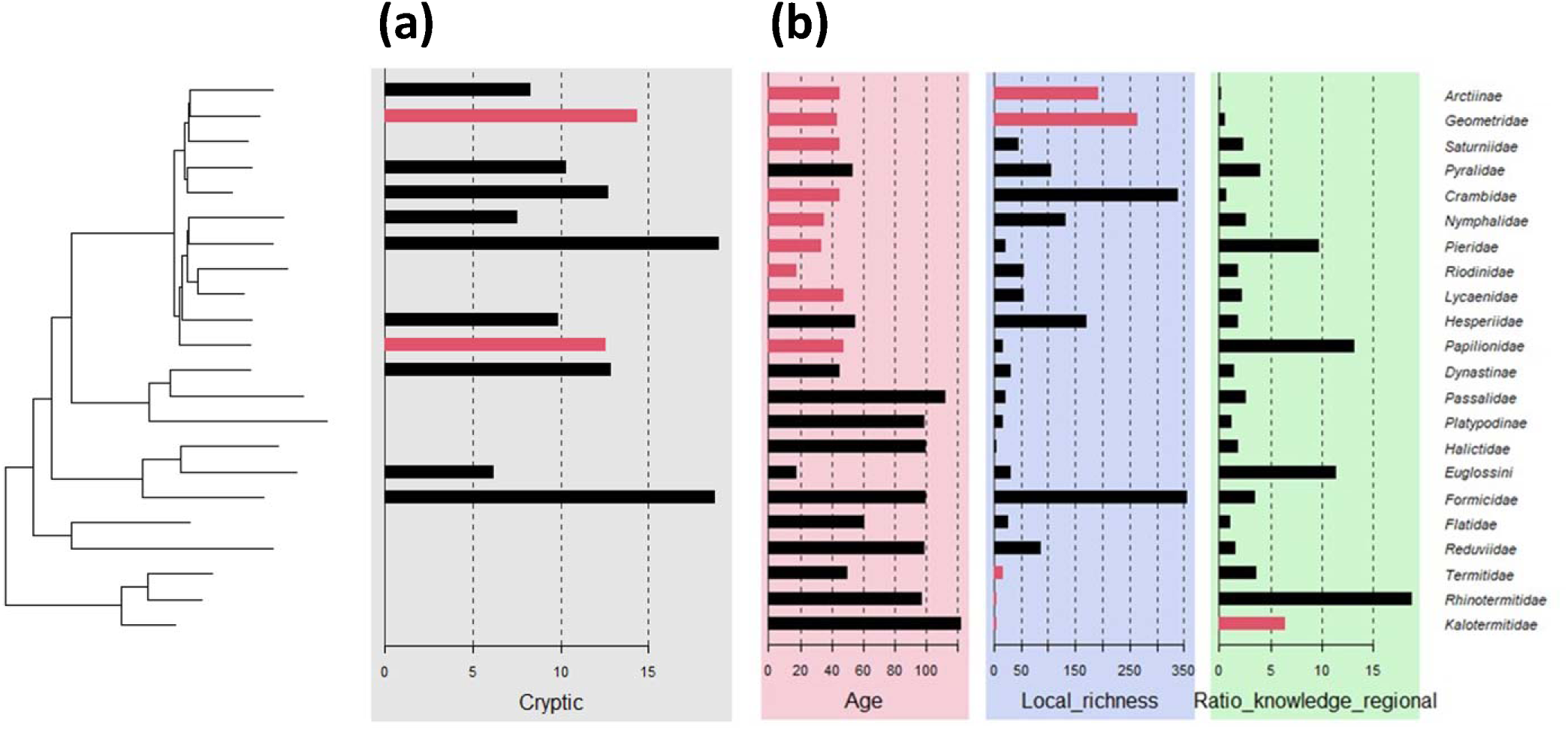
Local indicator of phylogenetic association (LIPA) based on Moran’s I for (a) percentage of cryptic species and (b) the three variables included in the best stepwise phylogenetic logistic regression. Colored bars indicate significant autocorrelation of trait values relative to their close relatives, i.e., significant phylogenetic signal for each tip.

## REFERENCES

Alfsnes, K., Leinaas, H.P. & Hessen, D.O. (2017) Genome size in arthropods; different roles of phylogeny, habitat and life history in insects and crustaceans. Ecology and Evolution, 7, 5939–5947.

Anderson-Teixeira, K.J., Davies S.J., Bennett A.C., Gonzalez-Akre E.B., Muller-Landau H.C., Wright S.J. et al. (2015) CTFS□Forest GEO: a worldwide network monitoring forests in an era of global change. Global Change Biology, 21, 528–549.

Bálint, M., Domisch, S., Engelhardt, C.H.M., Haase, P., Lehrian, S., Sauer, J. et al. (2011) Cryptic biodiversity loss linked to global climate change. Nature Climate Change, 1, 313–318.

Basset, Y., Barrios, H., Segar, S., Srygley, R.B., Aiello, A., Warren, A.D. et al. (2015) The butterflies of Barro Colorado Island, Panama: local extinction since the 1930s. PLoS One, 10, e0136623.

Basset, Y., Butterill, P.T., Donoso, D.A., Lamarre, G.P.A., Souto-Vilaros, D., Perez, F. et al. (2023) Abundance, occurrence and time series: long-term monitoring of social insects in a tropical rainforest. Ecological Indicators, 150, 110243.

Basset, Y., Eastwood, R., Sam, L., Lohman, D.J., Novotny, V., Treuer, T. et al. (2013) Cross-continental comparisons of butterfly assemblages in rainforests: implications for biological monitoring. Insect Conservation and Diversity, 6, 223–233.

Bastos□Silveira, C., Santos, S.M., Monarca, R., Mathias, M.D.L. & Heckel, G. (2012) Deep mitochondrial introgression and hybridization among ecologically divergent vole species. Molecular Ecology, 21, 5309–5323.

Beheregaray, L.B., & Caccone, A. (2007) Cryptic biodiversity in a changing world. Journal of Biology, 6, 1–5.

Belfiore, N.M., Hoffman, F.G., Baker, R.J. & Dewoody, J.A. (2003) The use of nuclear and mitochondrial single nucleotide polymorphisms to identify cryptic species. Molecular Ecology, 12, 2011–2017.

Bickford, D., Lohman, D.J., Sodhi, N.S., Ng, P.K., Meier, R., Winker, K. et al. (2007) Cryptic species as a window on diversity and conservation. Trends in Ecology & Evolution, 22, 148–155.

Bujan, J., Yanoviak, S.P., & Kaspari, M. (2016) Desiccation resistance in tropical insects: causes and mechanisms underlying variability in a Panama ant community. Ecology and Evolution, 6, 6282–6291.

Burns, J.M., Janzen, D.H., Hajibabaei, M., Hallwachs, W. & Hebert, P.D. (2008) DNA barcodes and cryptic species of skipper butterflies in the genus *Perichares* in Area de Conservacion Guanacaste, Costa Rica. Proceedings of the National Academy of Sciences, 105, 6350–6355.

Cheng, R., Luo, A., Orr, M., Ge, D., Hou, Z.E., Qu, Y. et al. (2024) Cryptic diversity begets challenges and opportunities in biodiversity research. Integrative Zoology, 20, 33–49.

Cheng, Z., Li, Q., Deng, J., Liu, Q. & Huang, X. (2023) The devil is in the details: Problems in DNA barcoding practices indicated by systematic evaluation of insect barcodes. Frontiers in Ecology and Evolution, 11, 1149839.

Chesters, D. (2020) The phylogeny of insects in the data□driven era. Systematic Entomology, 45, 540–551.

Cong, Y., Ye, X., Mei, Y., He, K. & Li, F. (2022) Transposons and non-coding regions drive the intrafamily differences of genome size in insects. IScience, 25, 104873.

D’Ercole, J., Dincă, V., Opler, P. A., Kondla, N., Schmidt, C., Phillips, J. D. et al. (2021) A DNA barcode library for the butterflies of North America. PeerJ, 9, e11157.

Dincă, V., Montagud, S., Talavera, G., Hernández-Roldán, J., Munguira, M.L., García-Barros, E. et al. (2015) DNA barcode reference library for Iberian butterflies enables a continental-scale preview of potential cryptic diversity. Scientific Reports, 5, 12395.

Ficetola, G.F. & Taberlet, P. (2023) Towards exhaustive community ecology via DNA metabarcoding. Molecular Ecology, 32, 6320–6329.

Fišer, C., Robinson, C.T. & Malard, F. (2018) Cryptic species as a window into the paradigm shift of the species concept. Molecular Ecology, 27, 613–635.

Hajibabaei, M., Janzen, D.H., Burns, J.M., Hallwachs, W. & Hebert, P.D. (2006) DNA barcodes distinguish species of tropical Lepidoptera. Proceedings of the National Academy of Sciences, 103, 968–971.

Hawksworth, D.L. (2012) Global species numbers of fungi: are tropical studies and molecular approaches contributing to a more robust estimate? Biodiversity and Conservation, 21, 2425–2433.

Hebert, P.D., Cywinska, A., Ball, S.L. & deWaard, J.R. (2003) Biological identifications through DNA barcodes. *Proceedings of the Royal Society of London*, Series B, 270, 313–321.

Hebert, P.D., Penton, E.H., Burns, J.M., Janzen, D.H. & Hallwachs, W. (2004) Ten species in one: DNA barcoding reveals cryptic species in the neotropical skipper butterfly *Astraptes fulgerator*. Proceedings of the National Academy of Sciences, 101, 14812–14817.

Hendrichs, J., Vera, T., De Meyer, M. & Clarke, A. (2015) Resolving cryptic species complexes of major tephritid pests. ZooKeys, 540: 5–39.

Hinojosa, J.C., Koubínová, D., Szenteczki, M.A., Pitteloud, C., Dincă, V., Alvarez, N. & Vila, R. (2019) A mirage of cryptic species: genomics uncover striking mitonuclear discordance in the butterfly *Thymelicus sylvestris*. Molecular Ecology, 28, 3857–3868.

Ho, L.S.T. & Ane, C. (2014) A linear-time algorithm for Gaussian and non-Gaussian trait evolution models Systematic Biology, 63, 397–408.

Honeycutt, R.L. (2021) DNA Barcodes: Controversies, Mechanisms, and Future Applications. Frontiers in Ecology and Evolution, 9, 718865.

Hsieh, T.C., Ma, K.H. & Chao, A. (2020) iNEXT: iNterpolation and EXTrapolation for species diversity. R package version 2.0.20 URL: http://chao.stat.nthu.edu.tw/wordpress/software-download/.

Ives, A.R. & Garland, T. Jr. (2010) Phylogenetic logistic regression for binary dependent variables. Systematic Biology, 59, 9–26.

Janzen, D.H., Hallwachs, W., Harvey, D.J., Darrow, K., Rougerie, R., Hajibabaei, M. et al. (2012) What happens to the traditional taxonomy when a well-known tropical saturniid moth fauna is DNA barcoded? Invertebrate Systematics, 26, 478–505.

Johnson, C.G. (1966) A functional system of adaptative dispersal by flight. Annual Review of Entomology, 11, 233–260.

Jones, G. (1997) Acoustic signals and speciation: the roles of natural and sexual selection in the evolution of cryptic species. Advances in the Study of Behaviour, 26, 317–354.

Jörger, K.M. & Schrödl, M. (2013) How to describe a cryptic species? Practical challenges of molecular taxonomy. Frontiers in Zoology, 10, 59.

Karanovic, T., Djurakic, M. & Eberhard, S.M. (2016) Cryptic species or inadequate taxonomy? Implementation of 2D geometric morphometrics based on integumental organs as landmarks for delimitation and description of copepod taxa. Systematic Biology, 65, 304–327.

Keck F., Rimet F., Bouchez A. & Franc A. (2016) phylosignal: an R package to measure, test, and explore the phylogenetic signal. Ecology and Evolution, 6, 2774–2780.

Lamarre, G.P.A., Fayle T.M., Segar, S.T., Laird-Hopkins, B., Nakamura, A., Souto-Vilarós, D. et al. (2020) Monitoring tropical insects in the 21st century. Advances in Ecological Research, 62, 295–330.

Leather, S.R. (2009) Taxonomic chauvinism threatens the future of entomology. Biologist, 56, 10–13.

Li, T.C., Wang, Y.K., Sui, Z.X., Wang, T., Nian, J.Z., Jiang, J.Z. et al. (2021) Multiple mitochondrial haplotypes within individual specimens may interfere with species identification and biodiversity estimation by DNA barcoding and metabarcoding in fig wasps. Systematic Entomology, 46, 887–899.

Li, X. & Wiens, J.J. (2023) Estimating global biodiversity: the role of cryptic insect species. Systematic Biology, 72, 391–403.

Lukhtanov, V.A., Dantchenko, A.V., Vishnevskaya, M.S. & Saifitdinova, A.F. (2015) Detecting cryptic species in sympatry and allopatry: analysis of hidden diversity in *Polyommatus* (*Agrodiaetus*) butterflies (Lepidoptera: Lycaenidae). Biological Journal of the Linnean Society, 116, 468–485.

Mayhew, P.J. (2007) Why are there so many insect species? Perspectives from fossils and phylogenies. Biological Reviews, 82, 425–454.

Meier, R., Blaimer, B.B., Buenaventura, E., Hartop, E., von Rintelen, T., Srivathsan, A. & Yeo, D. (2022) A re□analysis of the data in Sharkey et al.’s (2021) minimalist revision reveals that BINs do not deserve names, but BOLD Systems needs a stronger commitment to open science. Cladistics, 38, 264–275.

Miller, S.E. (2007) DNA barcoding and the renaissance of taxonomy. Proceedings of the National Academy of Sciences, 104, 4775–4776.

Murray, T.E., Fitzpatrick U., Brown, M.J.F. & Paxton, R.J. (2008) Cryptic species diversity in a widespread bumble bee complex revealed using mitochondrial DNA RFLPs. Conservation Genetics, 9, 653–666.

Nadler, S.A. & De León, G.P.P. (2011) Integrating molecular and morphological approaches for characterizing parasite cryptic species: implications for parasitology. Parasitology, 138, 1688–1709.

Núñez, R., Genaro, J.A., Pérez□Asso, A., Murillo□Ramos, L., Janzen, D.H., Hallwachs, W., et al. (2020) Species delimitation and evolutionary relationships among *Phoebis* New World sulphur butterflies (Lepidoptera, Pieridae, Coliadinae). Systematic Entomology, 45, 481–492.

Pfenninger, M. & Schwenk, K. (2007) Cryptic animal species are homogeneously distributed among taxa and biogeographical regions. BMC Evolutionary Biology, 7, 1–6.

Rainford, J.L., Hofreiter, M. & Mayhew, P.J. (2016) Phylogenetic analyses suggest that diversification and body size evolution are independent in insects. BMC Evolutionary Biology, 16, 1–17.

Ratnasingham, S. & Hebert, P.D. (2013) A DNA-based registry for all animal species: the Barcode Index Number (BIN) system. PloS one, 8, e66213.

Ribera, I., Barraclough, T.G. & Vogler, A.P. (2001) The effect of habitat type on speciation rates and range movements in aquatic beetles: inferences from species□level phylogenies. Molecular Ecology, 10, 721–735.

Schönrogge, K., Barr, B., Wardlaw, J.C., Napper, E., Gardner, M.G., Breen, J. et al. (2002) When rare species become endangered: cryptic speciation in myrmecophilous hoverflies. Biological Journal of the Linnean Society, 75, 291–300.

Smith, M.A., Woodley, N.E., Janzen, D.H., Hallwachs, W. & Hebert, P. D. (2006) DNA barcodes reveal cryptic host-specificity within the presumed polyphagous members of a genus of parasitoid flies (Diptera: Tachinidae). Proceedings of the National Academy of Sciences, 103, 3657–3662.

Stork, N.E. (2018) How many species of insects and other terrestrial arthropods are there on Earth? Annual Review of Entomology, 63, 31–45.

Struck, T.H., Feder, J.L., Bendiksby, M., Birkeland, S., Cerca, J., Gusarov, V.I. et al. (2018) Finding evolutionary processes hidden in cryptic species. Trends in Ecology & Evolution, 33, 153–163.

Srivathsan, A., Ang, Y., Heraty, J.M., Hwang, W.S., Jusoh, W.F., Kutty, S.N. et al. (2023) Convergence of dominance and neglect in flying insect diversity. Nature Ecology & Evolution, 7, 1012–1021.

Trontelj, P. & Fišer, C. (2009) Cryptic species diversity should not be trivialised. Systematics and Biodiversity, 7, 1–3.

Wahlberg, N. (2006) That awkward age for butterflies: insights from the age of the butterfly subfamily Nymphalinae (Lepidoptera: Nymphalidae). Systematic Biology, 55, 703–714.

Zhang, Q., Tong, X., Li, Y.Y., Sun, Q., Gao, Y., Zhang, S.H. et al. (2022) Presence of cryptic species in host insects forms a hierarchical Wolbachia infection pattern. Entomologia Generalis, 42, 571–578.

## References

D’Ercole, J., Dincă, V., Opler, P.A., Kondla, N., Schmidt, C., Phillips, J.D. et al. (2021) A DNA barcode library for the butterflies of North America. PeerJ, 9, e11157.

deWaard, J.R., Landry, J.F., Schmidt, B.C., Derhousoff, J., McLean, J.A. & Humble, L.M. (2009) In the dark in a large urban park: DNA barcodes illuminate cryptic and introduced moth species. Biodiversity and Conservation, 18, 3825–3839.

Gamboa, M., Muranyi, D., Kanmori, S. & Watanabe, K. (2019) Molecular phylogeny and diversification timing of the Nemouridae family (Insecta, Plecoptera) in the Japanese Archipelago. PloS one, 14, e0210269.

Hendrich, L., Morinière, J., Haszprunar, G., Hebert, P.D., Hausmann, A., Köhler, F. & Balke, M. (2015) A comprehensive DNA barcode database for Central European beetles with a focus on Germany: adding more than 3500 identified species to BOLD. Molecular Ecology Resources, 15, 795–818.

Janzen, D.H., Hallwachs, W., Blandin, P., Burns, J.M., Cadiou, J.M., Chacon, I. et al. (2009) Integration of DNA barcoding into an ongoing inventory of complex tropical biodiversity. Molecular Ecology Resources, 9, 1–26.

Kaartinen, R., Stone, G.N., Hearn, J., Lohse, K. & Roslin, T. (2010) Revealing secret liaisons: DNA barcoding changes our understanding of food webs. Ecological Entomology, 35, 623–638.

Kim, S., Lee, Y., Mutanen, M., Seung, J. & Lee, S. (2020) High functionality of DNA barcodes and revealed cases of cryptic diversity in Korean curved-horn moths (Lepidoptera: Gelechioidea). Scientific Reports, 10, 6208.

Lee, Y., Lee, W., Kanturski, M., Foottit, R.G., Akimoto, S.I. & Lee, S. (2017) Cryptic diversity of the subfamily Calaphidinae (Hemiptera: Aphididae) revealed by comprehensive DNA barcoding. PLoS one, 12, e0176582.

Lopez-Vaamonde, C., Kirichenko, N., Cama, A., Doorenweerd, C., Godfray, H.C.J., Guiguet, A. et al. (2021) Evaluating DNA barcoding for species identification and discovery in European gracillariid moths. Frontiers in Ecology and Evolution, 9, 626752.

Moraes, S.D.S., MurilloJRamos, L., Machado, P.A., Ghanavi, H.R., Magaldi, L.M., SilvaJBrandão, K.L. et al. (2021) A doubleJedged sword: Unrecognized cryptic diversity and taxonomic impediment in *Eois* (Lepidoptera, Geometridae). Zoologica Scripta, 50, 633–646.

Pedraza-Lara, C., Barrientos-Lozano, L., Rocha-Sánchez, A.Y. & Zaldívar-Riverón, A. (2015) Montane and coastal species diversification in the economically important Mexican grasshopper genus *Sphenarium* (Orthoptera: Pyrgomorphidae). Molecular Phylogenetics and Evolution, 84, 220–231.

Saito, R., Kato, S., Kuranishi, R.B., Nozaki, T., Fujino, T. & Tojo, K. (2018) Phylogeographic analyses of the *Stenopsyche* caddisflies (Trichoptera: Stenopsychidae) of the Asian Region. Freshwater Science, 37, 562–572.

Schmidt, S., SchmidJEgger, C., Morinière, J., Haszprunar, G. & Hebert, P.D. (2015) DNA barcoding largely supports 250 years of classical taxonomy: identifications for Central European bees (Hymenoptera, Apoidea partim). Molecular Ecology Resources, 15, 985–1000.

